# Gene expression profiling of skeletal muscles

**DOI:** 10.1101/2021.02.17.431599

**Authors:** Sarah I. Alto, Chih-Ning Chang, Kevin Brown, Chrissa Kioussi, Theresa M. Filtz

## Abstract

Soleus and tibialis anterior are two well-characterized skeletal muscles commonly utilized in skeletal muscle-related studies. Next-generation sequencing provides an opportunity for an in-depth biocomputational analysis to identify the gene expression patterns between soleus and tibialis anterior and analyze those genes’ functions based on past literature. This study acquired the gene expression profiles from soleus and tibialis anterior murine skeletal muscle biopsies via RNA-sequencing. Read counts were processed through edgeR’s differential gene expression analysis. Differentially expressed genes were filtered down using a false discovery rate less than 0.05c, a fold-change value larger than twenty, and an association with overrepresented pathways based on the Reactome pathway over-representation analysis tool. Most of the differentially expressed genes associated with soleus encoded for components of lipid metabolism and unique contractile elements. Differentially expressed genes associated with tibialis anterior encoded mostly for glucose and glycogen metabolic pathways’ regulatory enzymes and calcium-sensitive contractile components. These gene expression distinctions partly explain the genetic basis for muscle specialization and may help to explain skeletal muscle susceptibility to disease and drugs and refine tissue engineering approaches.

## Introduction

Six hundred individual skeletal muscles constitute 40% of a healthy adult human’s body mass. Skeletal muscle functions include, but are not limited to, voluntary body movement/locomotion, posture and body position, energy production, fatty acid (FA) β-oxidation, carbohydrate metabolism, and soft tissue support. Skeletal muscles consist of myofibers with defining metabolic and contractile properties.

In terms of metabolic properties, skeletal muscle is an important contributor to glucose, lipid, and protein metabolism. When mammals are in a healthy, well-fed state, skeletal muscle relies on carbohydrate or glucose metabolism as its primary means for energy. Adenosine triphosphate (ATP) can be generated from glucose for energy by aerobic respiration, anaerobic respiration, or both. Aerobic respiration produces a net total of 38 ATP molecules in four stages: glycolysis; a transition reaction that forms acetyl coenzyme A; the citric acid (Krebs) cycle; and an electron transport chain and chemiosmosis. Anaerobic respiration involves the transformation of glucose to lactate when oxygen is limited and produces only 2 ATP molecules, so it is only an effective means of energy production during a short period. Myofibers that utilize mostly aerobic respiration are called oxidative, those using primarily anaerobic respiration are called glycolytic, and ones that utilize both are called oxidative-glycolytic. Besides aerobic and anaerobic respiration, oxidative skeletal muscle can use FAs as a fuel source via β-oxidation. When consuming a high-fat diet or in a starving state, β-oxidation breaks down FAs, inhibits glucose metabolism, and decreases insulin sensitivity. As lipid resources become depleted, skeletal muscle proteins can be broken down for energy use.

Based on contractile properties, myofibers are defined by which myosin heavy chain is present. Myosin heavy chains (MyHCs) differentiate myofibers into types I (MYH7), II(A) (MYH2), II(B) (MYH4), and II(D or X) (MYH1) [1]. Myosin heavy chains are a significant component of the thick filament in a sarcomere. The heads of the myosin heavy chains move along the thin filament in the presence of ATP and calcium. The four contractile properties are connected with the three metabolic distinctions. Type I myofibers are oxidative [2]. Type II(A) myofibers can use anaerobic and aerobic respiration and are called oxidative-glycolytic [2]. Types II(B) and II(D) myofibers are characterized as glycolytic [2]. Type II(D) myofibers can combine with type II(A) or (B) to create hybrids called II(AD) and II(DB). Based on the metabolic and contractile properties, the myofiber types contract at specific rates starting at the slowest type I > II(A) > II(AD) > II(D) > II(DB) > II(B).

We studied two skeletal muscles from the mouse hindlimb that have a combination of the four myofiber types: Soleus (So) and Tibialis Anterior (Ta). The So and Ta muscles are antagonists to one another. When one muscle contracts (the agonist), the other relaxes (the antagonist), and vice versa. The So muscle is one of the superficial muscles that give shape to the lower hindlimb’s posterior component. So is essential for ankle joint movement to plantarflex the foot and maintain the standing posture [3]. The murine So is composed of approximately 50% type II(A), 40% type I, and 10% type II(D) myofibers [4]. Ta is located along the anterior lateral side of the tibia. It acts in the foot’s dorsiflexion and inversion, stabilizes the ankle when the foot hits the ground, and pulls the foot off the ground [3]. The murine Ta is composed of approximately 50-60% type II(B), 30-35% type II(D), and 5-10% II(A) myofibers [4].

The So and Ta muscles have been well-studied in numerous mouse studies that range from birth until postnatal day 28 (P28) and 3+ month-old adults. Mice become sexually mature between 4 to 8 weeks of age but continue to undergo rapid maturational growth until 3 to 6 months of age when they are mature adults. For instance, during development from P1 to P28, the murine Ta muscle increases seven-fold in fiber cross-sectional area and six-fold in mechanical function [5]. However, as an earlier study found, 8-week-old mice can handle more isometric stress than P28 mice [5,6]. Therefore, using P30 mice provides an opportunity to compare the myofibers’ metabolic and contractile properties in early adulthood.

In this study, differential gene expression profiling was examined in P30 murine So and Ta myofibers by RNA-seq. Sixteen molecular pathways were overrepresented among the differentially expressed (DE) genes between the So and Ta biopsies. The overrepresented pathways connect to glucose and glycogen metabolism, ion transport, and contraction. DE transcripts with a high-fold change that were not represented in the pathway analysis were further studied as well. These transcripts revealed molecular differences in the composition of neuromuscular junctions, immune system-related genes, and signaling pathways between the two muscle types. The detailed gene expression mapping of the two distinct skeletal muscles So and Ta will broaden our understanding of the skeletal muscles’ cellular diversity and potentially their differential susceptibility to myopathies and regenerating capabilities.

## Materials and methods

### Mice

All animal experiments were performed following the Institutional and National Health and Medical Research Council guidelines. The experimental protocol was approved by the Institutional Animal Care and Use Committee at Oregon State University. Biopsies from *Pitx2*^*FL/Z*^ mice were utilized as previously described for this study [7].

### RNA-Sequencing (RNA-Seq) data analysis

Six So myofiber biopsies (SRP127367, NCBI SRA) and two Ta myofiber biopsies (GSE114231, NCBI GEO, SRP145066, NCBI SRA) were collected at P30. RNA was extracted, sequenced, and analyzed as previously described [7]. The read counts were aligned with the mouse genome (mm10) to identify gene names with the read counts. TopHat2 was used to align and annotate over 24,000 RNA transcripts. The read counts were normalized to reduce transcript length bias [8]. Longer transcripts appear more prevalent and overshadow smaller transcripts if not corrected before processing differential expression analysis. The read counts were then processed using HTSeq [7]. Once aligned, normalized, and processed, The R package *edgeR* ran a statistical analysis to discover quantitative changes in gene expression levels between the So and Ta groups [9]. The *edgeR* program calculated the differential gene expression between genotypes based on a false discovery rate (FDR) cutoff of adjusted p-value < 0.05c and a log-fold change of +/− 1.0 [9]. Over 15,000 genes were identified as differentially expressed.

## Results

### RNA-Seq data quality check

This study compared the gene expression profiling in So and Ta skeletal muscles during the completion of sexual maturity in mice. Both muscles contain myofiber types with known differences in contractile and metabolic properties as well as different origins in development [1,2]. A comparison of differential gene expression between the muscles during this adolescent period has not been thoroughly explored.

Biopsies from So and Ta were processed and subject to RNA-Seq. The read counts were annotated into 24,421 genes using the mm10 genome reference. Gene expression comparisons revealed two separate clusters for So and Ta (Fig 1a) and identified 15,819 differentially expressed (DE) genes between the two muscles. All annotated DE genes were visualized in terms of log-base-2-fold change (log2(FC)) and the natural logarithm of the false discovery rate (Fig 1b). Of the 15,819 DE genes, 6,123 were statistically significant at an adjusted p-value < 0.05c and 2,481 had a greater than or equal to absolute 2-fold difference in expression between So and Ta. DE genes with at least one read count equal to zero were removed and normalized into z-scores (Fig 1c). The filtered dataset still retained separate groupings of myofiber from So and Ta.

**Fig 1.**
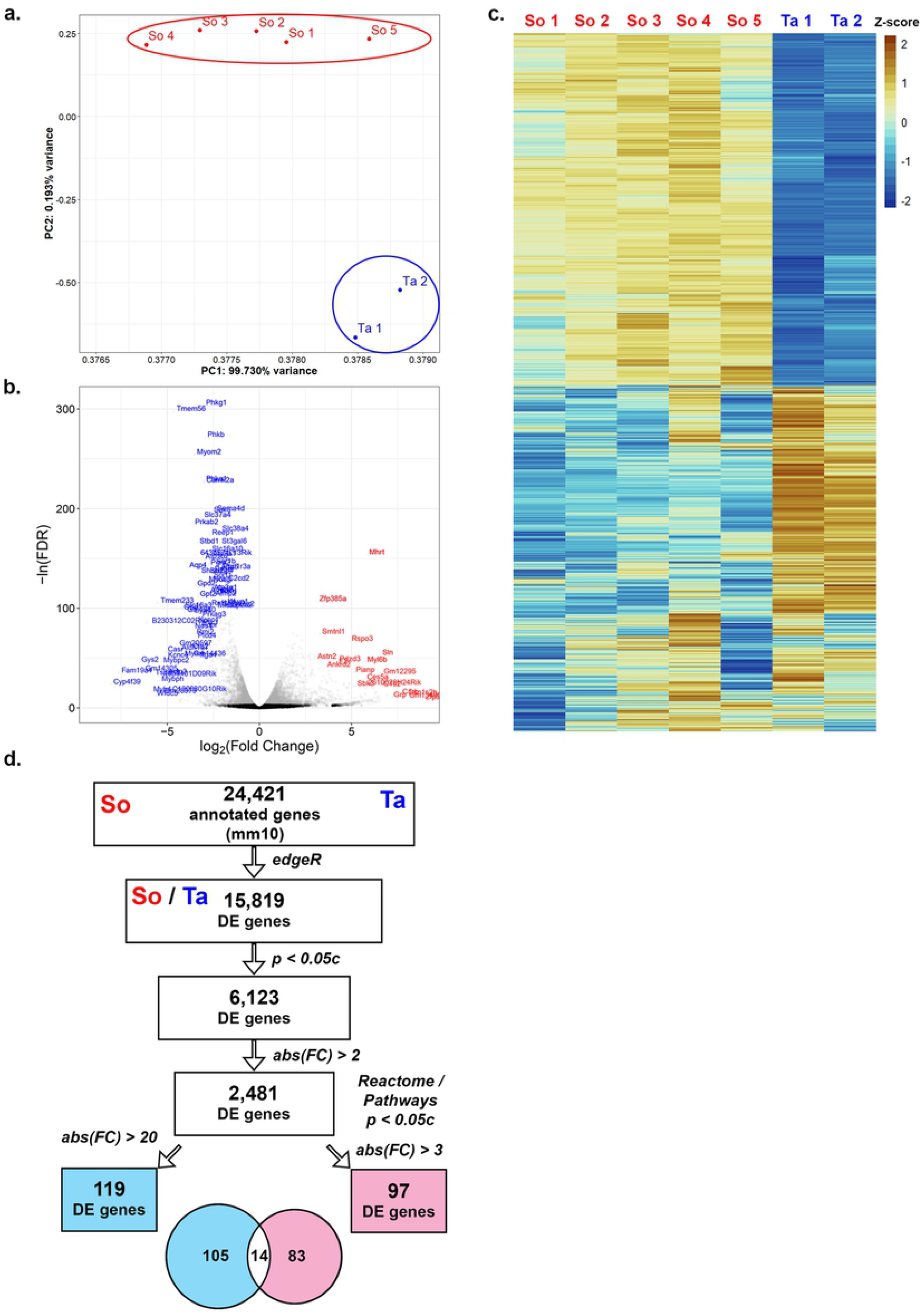
RNA-Seq workflow and data quality assessments. (A) Using the mm10 genome, 24,421 genes were annotated from soleus (So; n=5) and tibialis anterior (Ta; n=2) P30 biopsies. Total read counts from all annotated genes were subject to Principal Component Analysis (PCA) and plotted by sample groups from So (red font) or Ta (blue font). Throughout the analysis, increased gene expression from So was set to positive values, whereas negative values indicate decreased in So and, therefore, increased in Ta. (B) All DE genes were plotted with log2(FC) versus the −log transformation of false discovery rate. In the resultant volcano plot, DE genes with an adjusted p-value less than 0.05c are labeled as gray dots, and those with an adjusted p-value higher than 0.05c are black. Genes with increased expression in So or Ta are marked in red or blue font, respectively. (C) Read counts of genes were log-transformed and then normalized to z-scores with a mean of zero and standard deviation of one. Z-score values are graphically represented by the color intensity with z-scores above the mean in orange and z-scores below in blue. In the resultant heat map, columns represent each sample, and each row represents a statistically significant DE gene. (D) The annotated read counts were processed to determine differential expression between So and Ta using the R-package, *edgeR*. Sixteen molecular pathways were statistically overrepresented based on the Reactome database R package.

The molecular pathways associated with the DE genes were identified and statistically tested to determine if they were unique to myofibers. Using the Reactome database and its corresponding *ReactomePA* R package, the 2,481 DE genes were matched to molecular pathways, and those pathways were statistically analyzed for over- or under-representation [10,11]. The Reactome database employs Entrez ID numbers for gene identifications, so the number of genes analyzed by the Reactome R package decreased to 2,198 after removing genes with no Entrez ID. Hypergeometric testing statistically identified 19 Reactome pathways as overrepresented with an adjusted p-value of less than 0.05c. Three pathways had less than ten associated genes and were removed from further analysis. The DE genes related to the 16 Reactome pathways were ranked from highest to lowest log2(FC) values (S1 Fig). The ranking revealed that the 16 Reactome pathways could be consolidated into broader categories: contraction, ion & amino acid transport, and glucose & glycogen metabolism (S1 Fig).

The number of DE genes with greater than 3 FC and organized into the 16 Reactome-curated pathways, 97, (Fig 1d) was small compared to all the statistically significant DE genes from the input dataset (2481). Therefore, we reviewed the DE genes and found 119 with FC greater than or equal to 20 (high FC genes), of which only 14 were on the Reactome-curated list (Fig 1d). Manual examination of the genes’ known functions revealed that 36 of the remaining 105 high FC genes were easily placed into one of the three consolidated categories, demonstrating a limitation in the Reactome curation. An in-depth literature search was conducted for all 202 Reactome-curated and high FC DE genes to construct a clearer idea of the molecular differences between the So and Ta myofiber composition. Below, each consolidated category is addressed, highlighting some of the genes that may reveal or explain differences.

### Lipid metabolism

Skeletal muscles primarily metabolize glucose. Oxidative muscles can also use FAs along with glucose and glycogen as fuel sources. Free FA molecules are transported from the liver via cholesterol transport, which transports not only cholesterol and free FA molecules but also triglycerides and protein-bound FAs. Genes involved in cholesterol transport were present among the genes with the highest FC expression. Three cholesterol-related genes, *Pon1, Tspo2*, and *Apoa2*, were expressed 68, 51, and 27-fold higher in So, respectively (Fig 2, Table 1). *Pon1* protects against lipid oxidation for high-density lipoprotein (HDL) cholesterol and low-density lipoprotein (LDL) cholesterol [12]. *Tspo2* traffics free cholesterol in erythroid cells [13]. *Apoa2* regulates steroid concentrations by modulating cholesterol transport [14], the precursor of steroid hormones.

**Fig 2.**
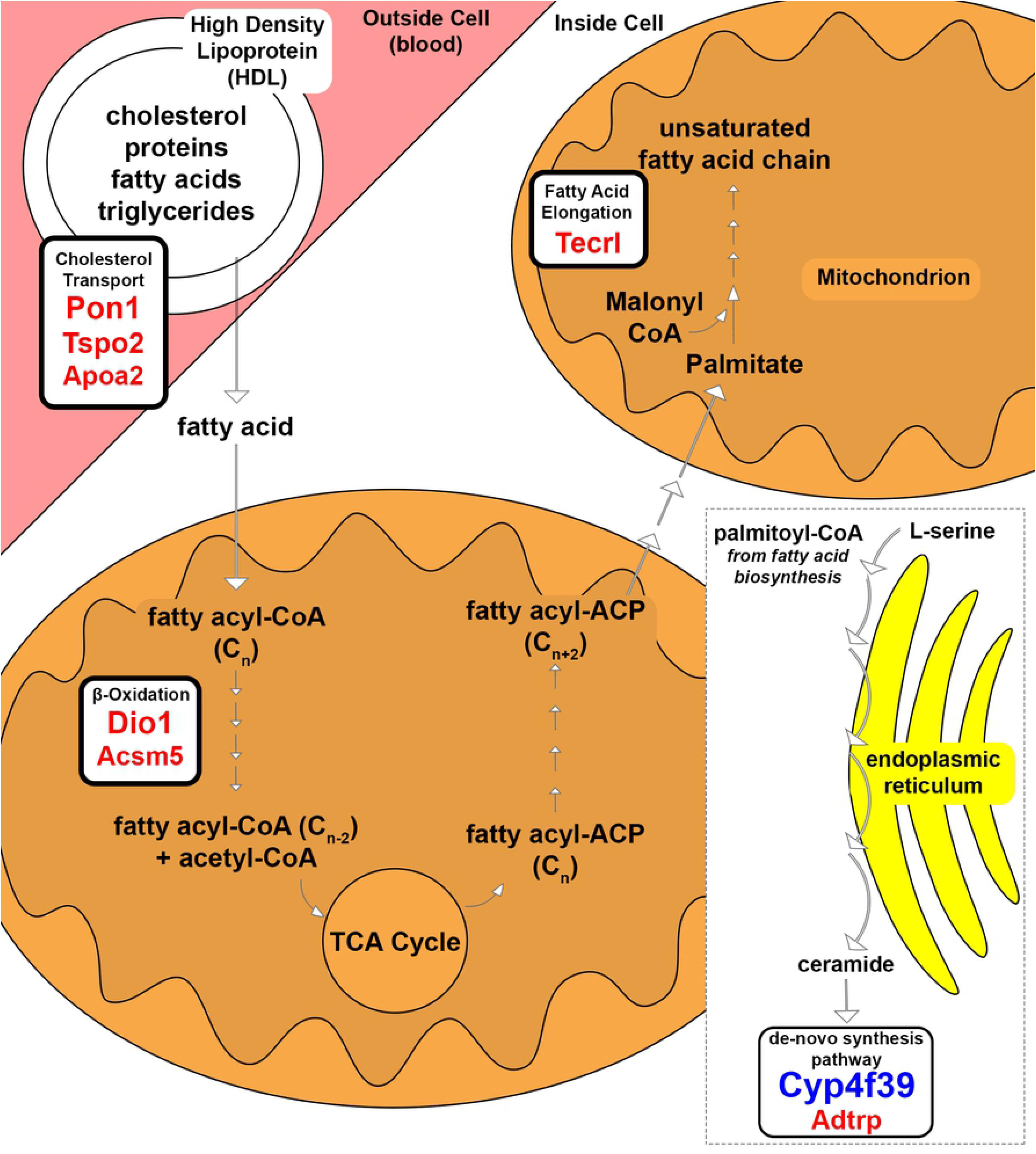
Lipid metabolism associated genes in So and Ta myofibers. Substrates and products of the enzymatic pathways for cholesterol transport, β-oxidation, FA elongation, and ceramide *de-novo* synthesis are shown based on their cellular location. The gene transcripts involved an enzymatic process and had increased expression in So or Ta are labeled in red or blue font, respectively. Font size correlates to relative FC.

**Table 1.** Differentially expressed genes associated with lipid, amino acid, and one-carbon metabolic pathways.

| Gene Symbol | Gene Name | Fold Change (So/Ta) | Localization and Biochemical Properties |
| --- | --- | --- | --- |
| **Metabolism** |  |  |  |
| **Lipid Metabolism** |  |  |  |
| *Dio1* | deiodinase, iodothyronine, type I | 69.1 | de-iodinate thyroid hormone interaction; FA oxidation; oxidative phosphorylation uncoupling; causing mitochondrial heat production [15] |
| *Pon1* | paraoxonase 1 | 68.83 | protection against oxidation for HDLs and LDLs [12] |
| *Tecrl* | trans-2,3-enoyl-CoA reductase-like | 55 | FA elongation in polyunsaturated FA biosynthesis [16] |
| *Tspo2* | translocator protein 2 | 51.12 | free cholesterol trafficking in erythroid cells [13] |
| *Acsm5* | acyl-CoA synthetase medium-chain family member 5 | 30.67 | FA β-oxidation [17] |
| *Apoa2* | apolipoprotein A-II | 27.64 | modulator of reverse cholesterol transport [14] |
| *Akr1c18* | aldo-keto reductase family 1, member C18 | 24.68 | catalyzes progesterone into 20-α-dihydroprogesterone (20-α-OHP) [18] |
| *Adtrp* | androgen dependent TFPI regulating protein | 21.66 | hydrolyzes FA esters of hydroxy FAs [19] |
| *Cyp4f39* | cytochrome P450, family 4, subfamily f, polypeptide 39 | -269.23 | 1ω and 2ω diesters types and cholesteryl (O-Acyl)-w-hydroxy FAs (OAHFAs) production; FA ω-hydroxylase; acyl-ceramide production [20] |
| **Amino acid Metabolism** |  |  |  |
| *Aox3* | aldehyde oxidase 3 | 30.48 | potentially linked to amino acid and retinol metabolisms; oxidizes aliphatic or aromatic aldehydes into carboxylic acid [21] |
| **One-Carbon Metabolism** |  |  |  |
| *Gnmt* | glycine N-methyltransferase | 26.87 | Methylation of DNA, RNA, proteins, and lipids via the methionine cycle |
| **Miscellaneous** |  |  |  |
| *Aldh1a7<sup>#</sup>* | aldehyde dehydrogenase family 1, subfamily A7 | -20.59 | Oxidoreductase, NAD/NADP acceptor [22] |

Positive or negative fold change represents increased gene expression in So or Ta, respectively. <sup>#</sup>DE gene both Reactome and 20x fold change

Once free FA molecules pass the plasma membrane, they enter the mitochondrial matrix to be broken down through β-oxidation. Specific protein channels handle different FA aliphatic tail lengths ranging from greater than 22 to fewer than 5 carbons. *Acsm5* was expressed 30 times higher in So, among the highest differentially expressed genes (Fig 2, Table 1). *Acsm5* encodes for an enzyme that catalyzes the activation of FAs with aliphatic tails of 6 to 12 carbons by CoA to produce acyl-CoA, the first step in FA metabolism [17].

Another gene, *Dio1*, was expressed 69-fold higher in So (Fig 2, Table 1). *Dio1* is involved in FA oxidation and oxidative phosphorylation uncoupling primarily in the liver, kidney, and thyroid [23] in addition to altering the thyroid hormone balance of triiodothyronine (T3) and thyroxine (T4) [15]. Thyroid hormone signaling regulates numerous genes involved in skeletal muscle homeostasis, function, metabolism [24].

Several of the highest FC DE genes in So are involved in the biosynthesis of lipids, sphingolipids, and derived lipids. FA molecules can be elongated and modified into complex and derived lipid molecules. *Tecrl*, involved in FA elongation in polyunsaturated FA biosynthesis [16] and myoblast differentiation upon TNF activation [25], was expressed 55-fold higher in So (Fig 2, Table 1). The increased expression of *Tecrl* could indicate that the So muscle is still developing in P30 mice.

Genes involved in pathways downstream of FA biosynthesis were also present among the study’s highest expressed genes. One of the highest FC genes in this study at 269-fold higher expression in Ta was *Cyp4f39*, which encodes for a FA ω-hydroxylase involved in acyl-ceramide synthesis [20] (Fig 2, Table 1). *Cyp4f39* is involved in cell surface protection, cell recognition, signaling, membrane transportation, increased rigidity, hydrolyzation of very long-chain FAs, and steroid hormone synthesis [20]. Ceramides have been linked to insulin resistance in type II diabetes [26].

### Glycogen metabolism

Glycogen metabolism combines two inverse processes: glycogenesis and glycogenolysis. Glycogenesis is a process where glucose or glucose 6-phosphate molecules are converted into glycogen, using 1 ATP and 1 ATP equivalent to add one glucose molecule to glycogen (Fig 3a). Glycolytic fibers store glycogen as a source of energy, whereas oxidative fibers do not. Glycogen is tapped during times of nutritional insufficiency or rapid demand for muscle activity. The direction of glycogen metabolism cannot be determined from this study alone, only inferring potentially emphasized steps in either So or Ta. The glycogenesis-involved enzymes phosphoglucomutase (*Pgm2, Pgm2l1*) and glycogen synthase (*Gys2*) had increased expression in Ta (Fig 3a; Table 2). Phosphoglucomutase reversibly converts glucose 6-phosphate to glucose 1-phosphate. Increased expression of phosphoglucomutase could contribute to either glycogen metabolic processes.

**Fig 3.**
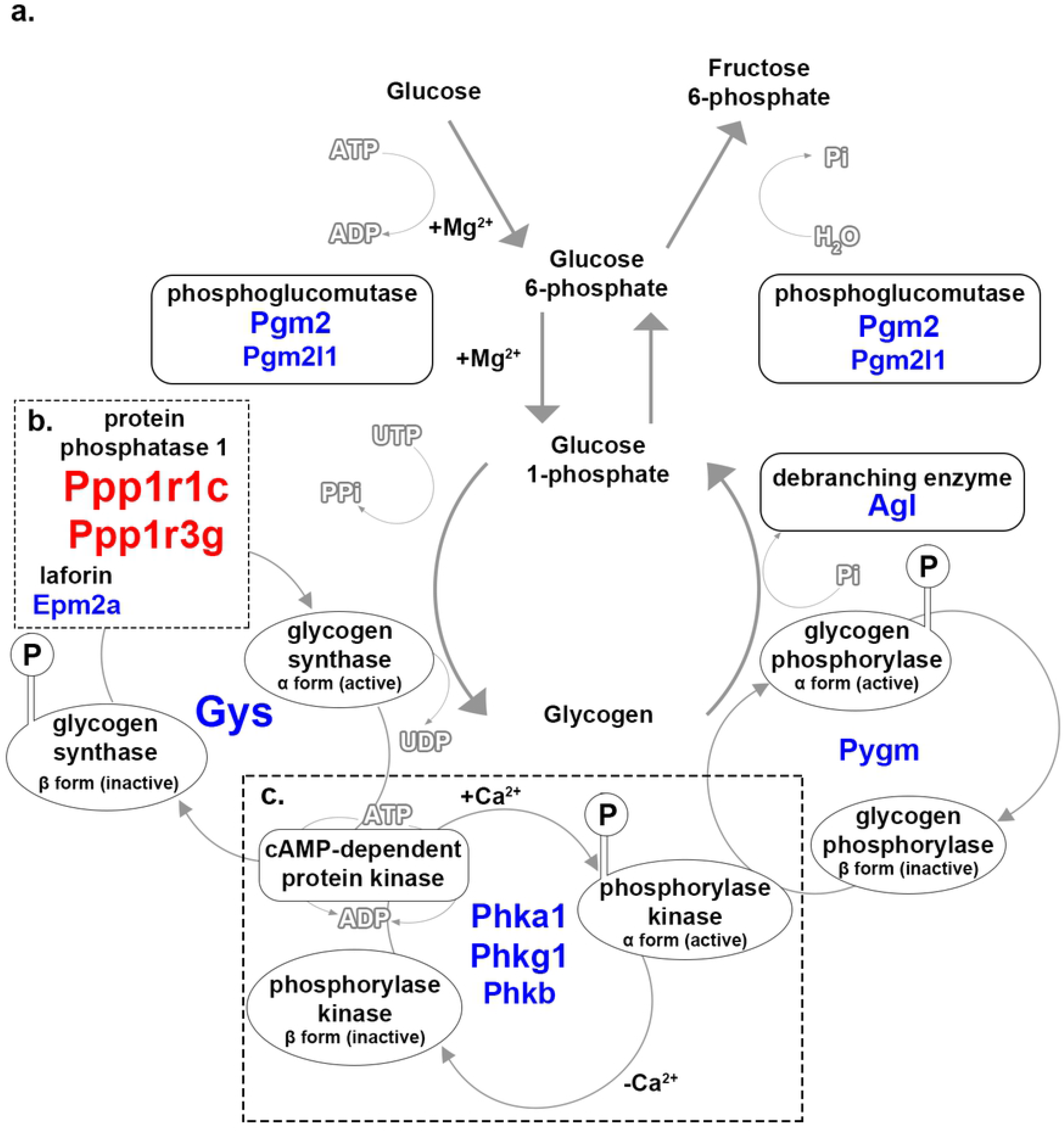
Glycogen metabolism associated genes in So and Ta myofibers. Substrates and products of the enzymatic steps in glycogen metabolism are represented, including the interconversion of inactive to active enzymatic states by regulatory enzymes. Enzymes involved at specific steps are encircled and adjacent to the corresponding reaction arrow. The gene symbols located underneath enzyme names in red or blue font were increased in So or Ta, respectively. Font size correlates to relative FC. (A) Glucose is converted into glycogen during glycogenesis. Glycogenolysis converts glycogen back into glucose molecules. Represented are magnesium ions (+Mg^2+^), adenosine diphosphate (ADP), uridine-5’-triphosphate (UTP), pyrophosphate (PPi), and inorganic phosphate (Pi). (B) Two subunits of the protein phosphorylase 1 had increased gene expression in So. The protein laforin had increased gene expression in Ta. Represented are an encircled ‘P’ indicating phosphorylation and uridine-diphosphate (UDP). (C) Three isoform subunits of phosphorylase kinase had increased gene expression in Ta. Represented are the addition of calcium ions (+Ca^2+^) and the removal of calcium ions (-Ca^2+^).

**Table 2.**
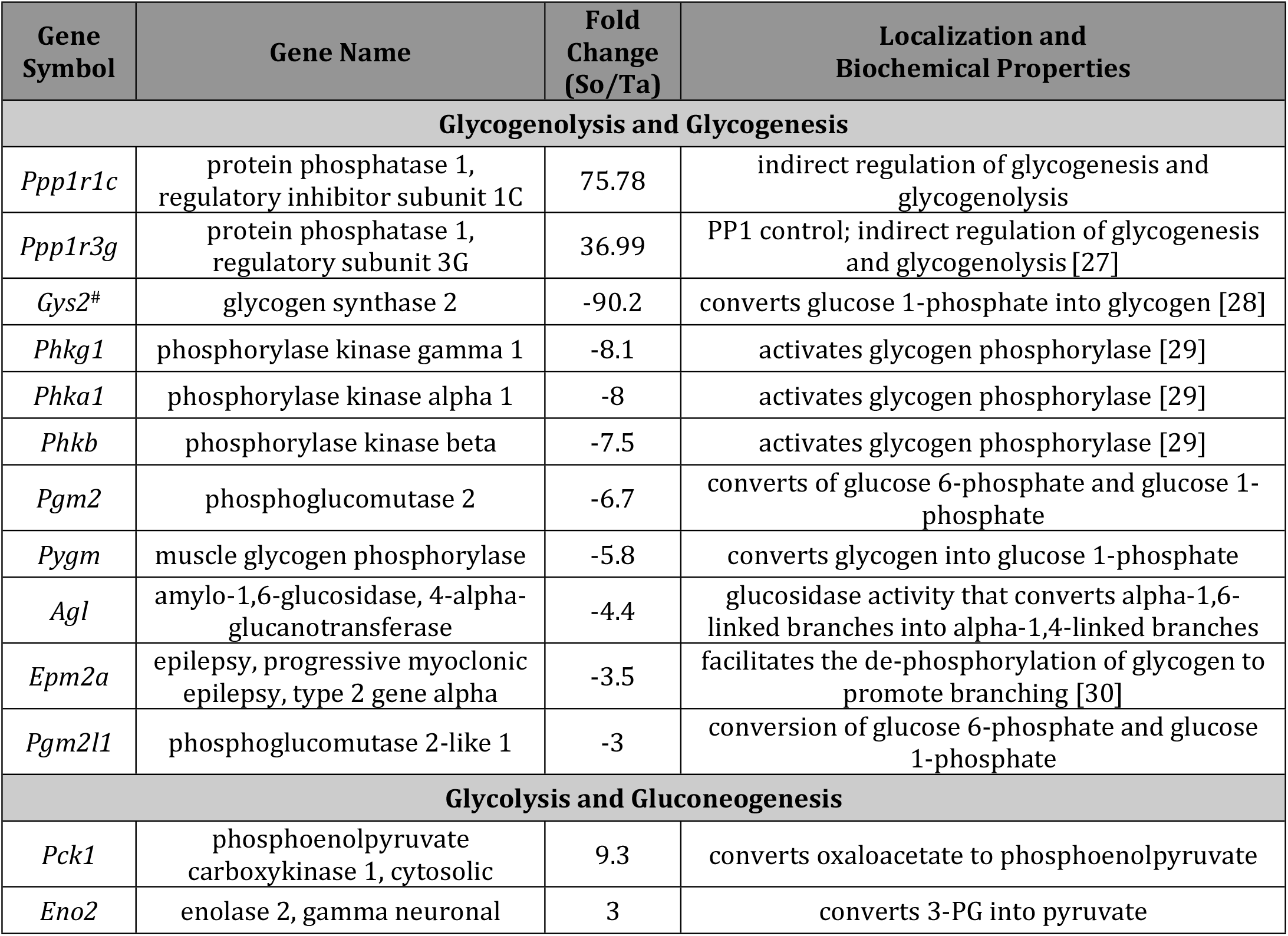
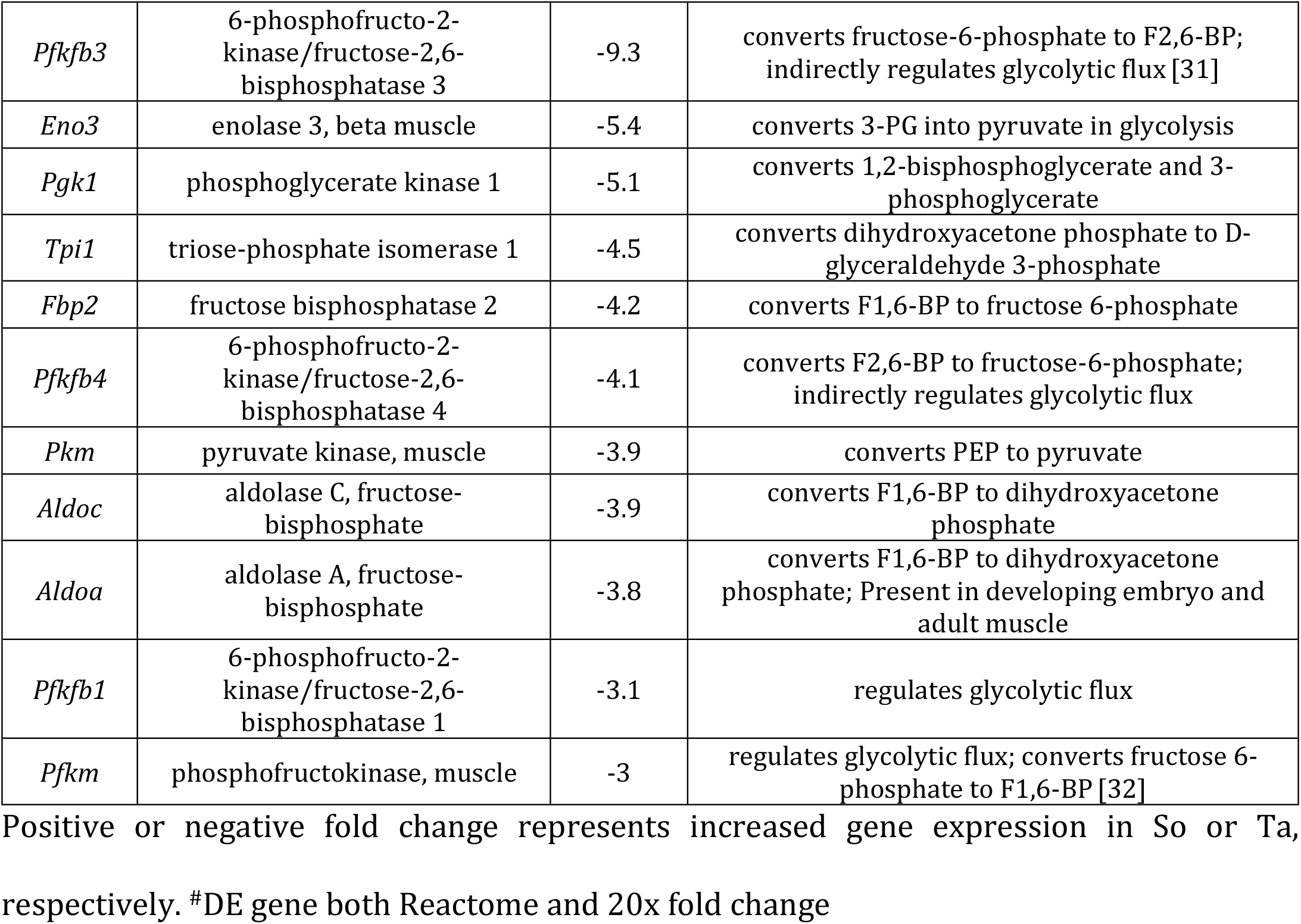
Differentially expressed genes associated with glucose and glycogen metabolic pathways.

Glycogen synthase is a critical enzyme in glycogenesis that converts glucose 1-phosphate into glycogen with the assistance of pyro-phosphorylase and a branching enzyme [27]. Increased gene expression of *Gys2* in Ta may imply glycogenesis is the priority in the Ta muscle (Fig 3a; Table 2). *Gys2* activity is regulated by dephosphorylation and phosphorylation by protein phosphatase 1 (PP1) and cyclic-adenosine monophosphate (AMP)-dependent protein kinase, respectively. PP1 is an enzyme complex with regulatory and catalytic subunits, of which two regulatory subunits, *Ppp1r1c* and *Ppp1r3g*, were expressed higher in So (Fig 3b; Table 2). The catalytic subunits of PP1 were present among the DE genes; however, they were not present in either the Reactome filtered or highest fold-change gene lists. *Ppp1r3g* regulates glycogen accumulation [28] and, along with the predicted inhibitory regulatory subunit *Ppp1r1c*, could be controlling glycogenesis in the So myofibers. The increased gene expression of the PP1 regulatory subunits in So could indicate a tighter control mechanism that prevents glycogen build-up. *Epm2a*, a factor that regulates PP1, was increased in Ta (Fig 3b; Table 2). *Epm2a* encodes for laforin, which interacts with PP1 and regulates the dephosphorylation of glycogen to promote glycogen branching [30]. Laforin is also involved in the prevention of cytotoxicity by protein ubiquitination [33] and it may control glycogen synthesis by ubiquitinating PP1, which would prevent activation of glycogen synthase.

Glycogenesis must be inhibited to trigger glycogenolysis or glycogen catabolism. Glycogenolysis neither requires nor produces energy when breaking glycogen into glucose molecules. The process is triggered by the accumulation of gluconeogenic precursors, such as lactate or alanine. Glycogen synthase is inactivated to inhibit glycogenesis. The inactivation triggers the activation of phosphorylase kinase, which phosphorylates glycogen phosphorylase. Phosphorylase kinase is an enzyme complex of four subunits [29], three of which (*Phka1, Phkg1, Phkb*) also had increased expression in Ta (Fig 3c; Table 2). Phosphorylase kinase could be regulating the glycogenolysis in Ta without inhibiting glycogen synthase by ubiquitination. Glycogen phosphorylase pairs with a debranching enzyme to break off glucose 1-phosphate molecules from glycogen; a gene encoding for each (*Pygm* and *Agl*, respectively) was expressed higher in Ta (Fig 3a; Table 2).

### Glucose metabolism

Glucose metabolism was one of the overrepresented molecular pathways identified. Glucose metabolism consolidates glycolysis and gluconeogenesis enzymatic processes. The former process breaks glucose into an intermediate, pyruvate, and the latter is the reverse process. Glycolysis converts glucose into pyruvate in 10 steps consuming one glucose, 2 NAD^+^, 2 Pi, 4 ADP, and 2 ATP molecules to generate 2 pyruvate, 2 NADH, 2 H^+^, 4 ADP, and 4 ATP molecules. The net products are 2 pyruvate, 2 ATP, 2 NADH, and 2H^+^ molecules. Gluconeogenesis converts 2 pyruvate, 4 ATP, 2 GTP, 2 NADH, 2 H^+^, and 2 H2O molecules into fructose 6-phosphate or glucose 6-phosphate. Overall, gluconeogenesis consumes 6 ATP, and glycolysis generates a net 2 ATP. Therefore, the whole glycolysis/gluconeogenesis cycle costs four ATP.

Three critical enzymes regulate the whole glycolysis process: hexokinase, phosphofructokinase-1 (PFK1), and pyruvate kinase. Genes encoding for PFK1 (*Pfkm*) and pyruvate kinase (*Pkm*) had increased expression in Ta (Fig 4a; blue font, Table 2). PFK1 converts fructose 6-phosphate to F1,6-BP and is activated by high AMP concentrations and fructose 2,6-bisphosphate (F2,6-BP) [32]. Pyruvate kinase converts phosphoenolpyruvate to pyruvate when activated by high F1,6-BP concentration and is deactivated by an elevated ATP concentration. Four other isoenzymes involved in glycolysis and gluconeogenesis were increased in Ta: aldolase (*Aldoa, Aldoc*), triosephosphate isomerase (*Tpi1*), phosphoglycerate kinase (*Pgk1*), and enolase (*Eno3* in Ta and *Eno2* in So) (Fig 4a, Table 2).

**Fig 4.**
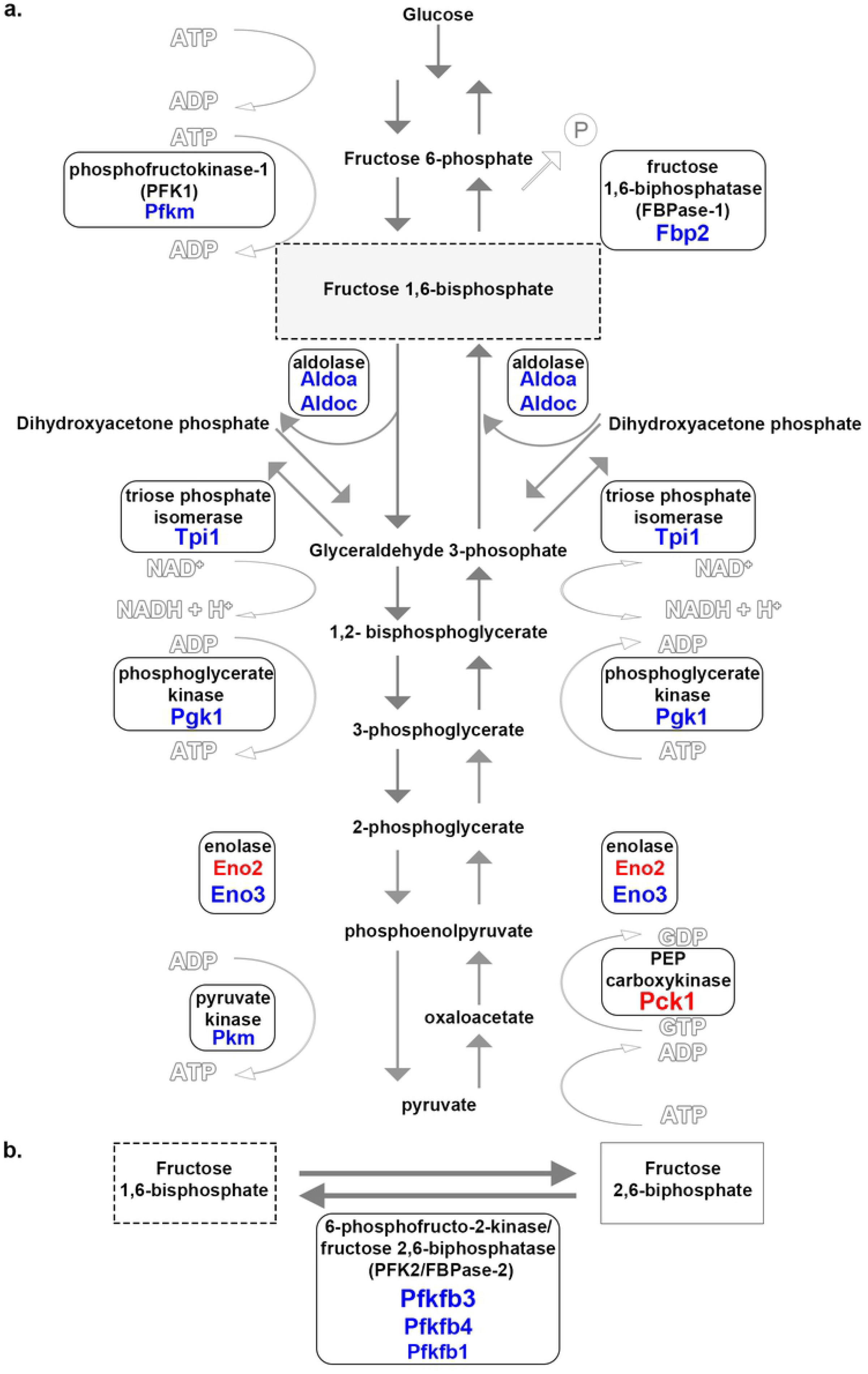
Glucose metabolism associated genes in So and Ta myofibers. Substrates and products of the enzymatic steps in glucose metabolism are represented. Enzymes involved at specific stages are encircled and adjacent to the corresponding reaction arrow. The red or blue font of gene symbols underneath enzyme names represent increased expression in So or Ta, respectively. Font size correlates to relative FC. (A) The isoenzymes encoded by *Aldoa, Aldoc, Tpi1, Pgk1, Eno3, Pfkm, Pkm*, and *Fbp2* had increased gene expression in Ta. Isoforms of enolase (*Eno2*) and PEP carboxykinase (*Pck1*) had increased gene expression in So. Represented are guanosine triphosphate (GTP), guanosine diphosphate (GDP), and inorganic phosphate (P). (B) Subunits of the PFK2/FBPase2 complex, *Pfkfb3, Pfkfb1*, and *Pfkfb4*, were increased in Ta.

The indirect regulator of glycolysis is called 6-phosphofructo-2-kinase/ fructose 2,6-bisphosphatase (PFK2/FBPase2) complex. The PFK2/FBPase2 complex pulls fructose 6-phosphate out of the glycolytic process and converts it to F2,6-BP [31]. In Ta, three isoenzymes of PFK2/FBPase2, *Pfkfb3, Pfkfb1*, and *Pfkfb4* had increased expression by 3.1, 4.1, and 9.3-fold, respectively (Fig 4b; blue font, Table 2).

When blood glucose concentrations are low, glucose can be generated through gluconeogenesis, an 11-step process for converting pyruvate to glucose. The enzymes pyruvate carboxylase, PEP carboxykinase, and fructose 1,6-bisphosphatase (FBPase1) are critical regulatory points in gluconeogenesis. FBPase1 (*Fbp2*) was increased in Ta, and PEP carboxykinase (*Pck1*) in So (Fig 4a, Table 2). FBPase1 converts F1,6-BP to fructose 6-phosphate in the presence of a high concentration of AMP and F2,6-BP, while increased citrate levels deactivate it. PEP carboxykinase turns oxaloacetate into phosphoenolpyruvate and is enabled by a high ADP concentration.

All myofibers utilize glycolysis, whether or not they use aerobic and/or anaerobic respiration. Glycolysis occurs under both aerobic and anaerobic conditions because it does not require oxygen. The majority of the DE genes associated with glucose metabolism had prominent expression in Ta compared to So. As a muscle that uses anaerobic metabolism to gain fuel, Ta may better utilize the anaerobic pathway due to the increased expression levels of glycolytic and regulatory enzymes.

### Contraction

The Reactome pathway overrepresentation analysis identified contraction as another significant difference between So and Ta. The molecular basis of muscle contraction is explained by the sliding filament theory, where myosin and actin filaments interact in a particular manner to produce contractile force. Most contraction-related DE genes exposed were associated with the thin filament, thick filament, and Z-line of the sarcomere (Fig 5a, Table 3).

**Fig 5.**
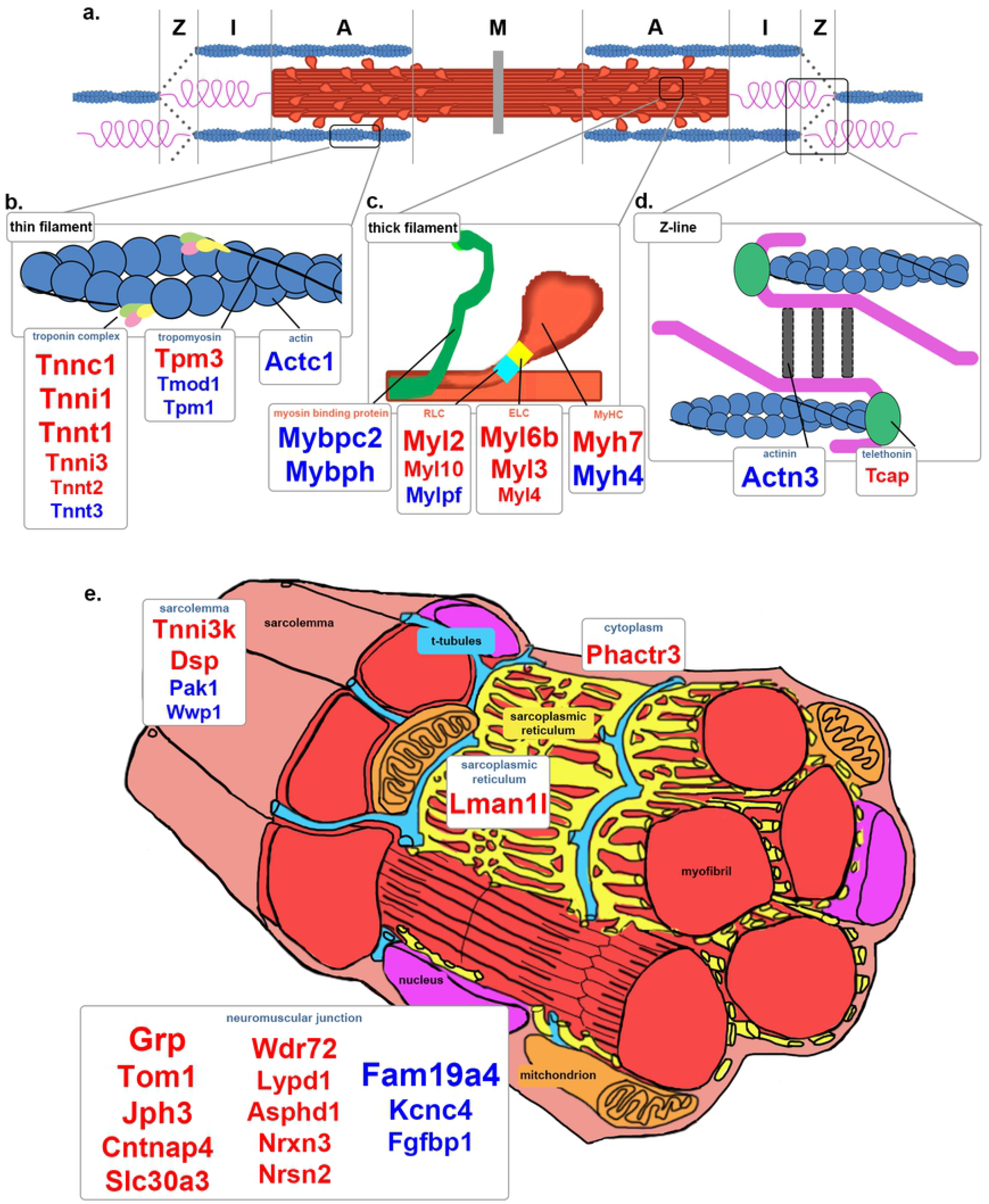
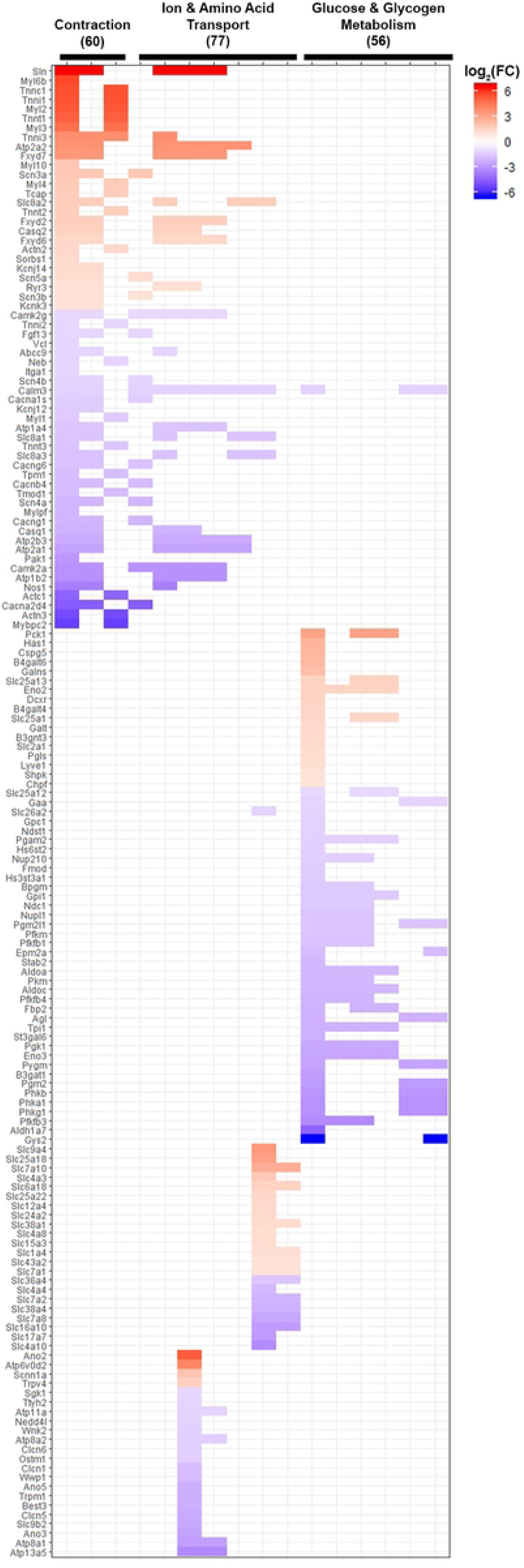
Contraction related genes in So and Ta. (A) *Graphic representation of the sarcomere*. The M-line, A-zone, I-zone, and Z-line are depicted. The red band represents the thick myosin filament. The blue spheres represent the thin filament. Within the I-zone also resides an elastic protein called titin. (B) *Graphic representation of thin filament structure*. The actin filament and other proteins associated with the thin filament are depicted here. *Tpm3, Tnnc1, Tnni1, Tnnt1, Tnni3, Tnnt2*, and *Tnnt3* were increased in So. *Actc1, Tmod1, Tpm1*, and *Tnnt3* were increased in Ta. (C) *Thick filament close-up*. The myosin heavy chain heads and other proteins aiding in myosin head movement are depicted here. *Myh7, Myl6b, Myl3, Myl4, Myl2*, and *Myl10* were increased in So. *Myh4, Mylpf, Mybpc2*, and *Mybph* were increased in Ta. (D) *Z-line close-up*. The Z-line region of the sarcomere is where actin and titin are connected to an antiparallel complex, including titin cap (*Tcap*) and actinin (*Actn3*). (E) *Myofiber structure*. Myofibers are multinucleated cells with extensive bundled networks of myofibrils, sarcoplasmic reticulum, t-tubules, and mitochondria. T-tubules transmits nerve impulses from neuromuscular junctions to myofibrils to stimulate contraction. *Jph3, Lman1l, Phactr3, Tnni3k, Dsp, Grp, Tom1, Cntnap4, Slc30a3, Wdr72, Lypd1, Asphd1, Nrxn3*, and *Nrsn2* were increased in So. *Pak1, Wwp1, Fam19a4, Kcnc4*, and *Fgfbp1* were increased in Ta. Red or blue gene symbols underneath structure names represent increased expression in So or Ta, respectively. Font size correlates to relative FC.

**Table 3.** Differentially expressed genes associated with contractile related structures.

| Gene Symbol | Gene Name | Fold Change (So/Ta) | Localization and Biochemical Properties |
| --- | --- | --- | --- |
| **Neuromuscular Junction** |  |  |  |
| *Grp* | gastrin releasing peptide | 149.61 | modulates autonomic system; regulates male sexual function; conveys itch sensation |
| *Tom1* | target of myb1 trafficking protein | 84.21 | recruits clathrin to endosomal structures [34] |
| *Jph3* | junctophilin 3 | 66.32 | Formation of junctional membrane complexes; motor coordination [35] |
| *Cntnap4* | contactin associated protein-like 4 | 58.74 | cell adhesion molecule and receptor [36] |
| *Slc30a3* (*Znt3*) | solute carrier family 30 (zinc transporter), member 3 | 34.8 | transport of zinc in synaptic vesicles |
| *Wdr72* | WD repeat domain 72 | 33.63 | endocytosis, protein reabsorption, and calcium excretion [37] |
| *Lypd1* | Ly6/Plaur domain containing 1 | 26.17 | regulates neuronal nicotinic receptors |
| *Asphd1* | aspartate β-hydroxylase domain-containing protein 1 | 25.7 | possibly neurotransmission and synaptic vesicle location/function |
| *Nrxn3* | neurexin III | 24.74 | synapse organization; regulating neurotransmitter release [38] |
| *Nrsn2* | neurensin 2 | 24.72 | transporting small vesicles to perinuclear region to exit towards organelles |
| *Fam19a4* (*Tafa4*) | TAFA chemokine like family member 4 | -199.82 | modulates neuronal excitability and controls somatic sensation threshold [39] |
| *Kcnc4* (*Kv3.4*) | potassium voltage gated channel, Shaw-related subfamily, member 4 | -33.68 | broadens action potential to prevent somatic depolarization [40] |
| *Fgfbp1* | fibroblast growth factor binding protein 1 | -28.02 | slows neuromuscular junctions (NMJs) degeneration [41] |
| **Sarcolemma, Sarcoplasm, and Sarcoplasmic Reticulum** |  |  |  |
| *Lman1l* | lectin, mannose-binding 1 like | 61.63 | binds to glycoproteins transported from endoplasmic reticulum (ER) to ER-Golgi intermediate [42] |
| *Tnni3k* | TNNI3 interacting kinase | 27.58 | promotes oxidative stress and myocyte death; part of costamere attached to sarcolemma [43] |
| *Phactr3* | phosphatase and actin regulator 3 | 26.09 | binds to cytoplasmic actin and regulates PP1 |
| *Dsp* | desmoplakin | 20.92 | assembles functional desmosomes; maintains cytoskeletal architecture [44] |
| *Pak1* | p21 (RAC1) activated kinase 1 | -7.3 | regulates muscle-specific kinase to maintain NMJs, actin remodeling, and glucose uptakes in skeletal muscle |
| *Wwp1* | WW domain containing E3 ubiquitin protein ligase 1 | -3.5 | cell proliferation and apoptosis |
| **Thin Filament** |  |  |  |
| **Actin** |  |  |  |
| *Actc1<sup>#</sup>* | actin, alpha, cardiac muscle 1 | -20.7 | Assembles into filaments that are involved in muscle contraction, cell motility, cell signaling, & vesicle movement; associated with fetal skeletal muscle |
| **Tropomyosin & Tropomodulin** |  |  |  |
| *Tpm3* | tropomyosin 3, gamma | 29.17 | controls the actin filament with tropomodulin [45] |
| *Tmod1* | tropomodulin 1 | -3.6 | controls the Ca<sup>2+</sup>-regulated thin filament end with tropomyosin |
| *Tpm1* | tropomyosin 1, alpha | -3.4 | controls the Ca<sup>2+</sup>-regulated thin filament end with tropomodulin [46] |
| **Troponin Complex** |  |  |  |
| *Tnnc1<sup>#</sup>* | troponin C, cardiac/slow skeletal | 47.7 | binds to calcium and exposes the myosin head binding sites; slow skeletal muscle |
| *Tnni1<sup>#</sup>* | troponin I, skeletal, slow 1 | 46.2 | binds to actin and inhibits ATPase activity; slow skeletal muscle |
| *Tnnt1<sup>#</sup>* | troponin T1, skeletal, slow | 36.6 | anchors to tropomodulin; slow skeletal muscle [47] |
| *Tnni3* | troponin I, cardiac 3 | 14.3 | binds to actin and inhibits ATPase activity; cardiac muscle |
| *Tnnt2* | troponin T2, cardiac | 3.3 | anchors to tropomodulin; cardiac, embryonic, and neonatal skeletal muscles [48] |
| *Tnnt3* | troponin T3, skeletal, fast | -3 | anchors to the tropomodulin; fast skeletal muscle [49] |
| **Thick Filament** |  |  |  |
| **Myosin Heavy Chains** |  |  |  |
| *Myh7* | myosin, heavy polypeptide 7, cardiac muscle, beta | 34.71 | Isoform of myosin present in slow (type I) skeletal muscle fibers [1] |
| *Myh4* | myosin, heavy polypeptide 4, skeletal muscle | -57.73 | Isoform of myosin present in adult type IIB skeletal muscle fibers [50] |
| **Essential Myosin Light Chains** |  |  |  |
| *Myl6b<sup>#</sup>* | myosin, light polypeptide 6B | 54.1 | controls cell adhesion, cell migration, tissue architecture, cargo transport, and endocytosis; promotes p53 protein ubiquitination and degradation |
| *Myl3*<sup>#</sup> | myosin, light polypeptide 3 | 27 | involved in force development and fine-scale coordinated muscle contraction |
| *Myl4* | myosin, light polypeptide 4 | 3.6 | Binds to Ca<sup>2+</sup>; embryonic skeletal muscle and atrial myocardium [2] |
| **Regulatory Myosin Light Chains** |  |  |  |
| *Myl2*<sup>#</sup> | myosin, light polypeptide 2, regulatory, cardiac, slow | 41.7 | stiffens myosin neck and aids in myosin head movement |
| *Myl10* | myosin, light chain 10, regulatory | 4 | stiffens myosin neck and aids in myosin head movement [51] |
| *Mylpf* | myosin light chain, phosphorylatable, fast skeletal muscle | -4 | regulates myofilament activation by phosphorylation [52] |
| **Myosin Binding Proteins** |  |  |  |
| *Mybpc2*<sup>#</sup> | myosin binding protein C, fast type | -40.7 | regulates of myofilament contraction [53] |
| *Mybph* | myosin binding protein H | -42.77 | sarcomere contraction; maturation process of auto-phagosomal membranes; inhibition of non-muscle RLC MYL2A and MYH2A |
| **Z-line** |  |  |  |
| *Tcap* | titin-cap | 3.5 | binds to titin in an anti-parallel complex and stabilizes Z-line [54] |
| *Actn3*<sup>#</sup> | actinin alpha 3 | -33.3 | anchors actin filaments and supports sarcomere [55] |

Positive or negative fold change represents increased gene expression in So or Ta, respectively. <sup>#</sup>DE gene both Reactome and 20x fold change lists

The thin filament is made up of actin subunits. *Actc1* had increased expression in Ta (Fig 5b, Table 3). Tropomyosin wraps outside of the actin helix to stabilize the strand and rotate the thin filament to expose the binding sites. Two isoforms of tropomyosin had increased gene expression in So (*Tpm3*) [45] or Ta (*Tpm1*) [46] (Fig 5b, Table 3). The troponin complex is a regulatory element of the actin filament and is associated with calcium binding, inhibitory regulation, and tropomyosin binding. Tropomyosin-binding element encoding genes *Tnnt1* [47] and *Tnnt2* [48] were increased in So while Ta had increased expression of *Tnnt3* [49] (Fig 5b, Table 3). The calcium and inhibitory units *Tnnc1, Tnni1*, and *Tnni3* had an increased expression in So (Fig 5b, Table 3). The actin filaments are capped by tropomodulin *Tmod1* expressed higher in Ta (Fig 5b, Table 3).

The thick filament consists of several strands of heavy myosin chains. Two myosin heavy chains wrapped around one another with myosin heads at the ends that attach and flex to move along the thin filament. Two myosin heavy chain isoforms unique to type I (*Myh7*) [1] and type II(B) (*Myh4*) [50] myofibers had increased gene expressions in So and Ta, respectively (Fig 5c, Table 3).

Myosin light chains and myosin-binding proteins regulate myosin head movement. Myosin light chains are classified into essential (ELCs) and regulatory (RLCs). ELCs stabilize the myosin head while RLCs stiffen the myosin neck domain. ELCs encoded by *Myl6b, Myl3*, and *Myl4* [2] and the RLCs encoded by *Myl2, Myl10* [51] were increased in So (Fig 5c, Table 3) while only *Mylpf* [52] was increased in Ta. *Mybpc2* [53] and *Mybph* that encode for myosin-binding proteins that assist in movement along the thin filament were increased in Ta (Fig 5c, Table 3).

The thin and thick filaments are held in an antiparallel configuration by a structure called the Z-line at the sarcomere ends (Fig 5d). The titin cap (*Tcap*) [54] is involved in mechano-electrical links between Z-lines and T-tubules and was expressed higher in So (Fig 5d, Table 3). Actinin (*Actn3*) [55] is involved in stabilizing the antiparallel formation and was increased in Ta (Fig 5d, Table 3). These structural differences could be altering the electrical activity and, in turn, the contractile properties of the different myofibers in So and Ta muscles.

Several of the highest FC DE genes were part of the neuromuscular junction (NMJ) apparatus. Several genes associated with the presynaptic membrane side, *Grp, Slc30a3, Asphd1, Nrxn3* [38], and *Nrsn2*, were increased in So, while *Fam19a4* [39], *Kcnc4* [40], and *Fgfbp1* [41] were increased in Ta (Fig 5e, Table 3). On the postsynaptic membrane side, *Tom1* [34], *Cntnap4* [36], *Wdr72* [37], and *Lypd1* were increased in So (Fig 5e; red font, Table 3). Several of these genes are associated with endocytosis and neurotransmitter receptor signaling.

## Discussion

Two broad-scale patterns emerged along with the three categories of overrepresented molecular pathways when examining the differential gene expression profiles of So and Ta muscles (S1 Fig). First, DE genes involved in lipid, glucose, and glycogen metabolism pathways associated more with one muscle than the other (Tables 1 and 2). Second, each muscle expressed unique genes encoding for similarly functioning isoenzymes, particularly for genes involved in the contraction and ion transport categories (Table 3 and S1 Table). We put these differences into context below, starting with the pathways that differentiate the muscle types. DE genes with the highest FC difference were not prevalent among genes associated with the overrepresented pathways. Therefore, DE genes with a greater than 20-fold expression difference were included for further investigation. Similarly seen in a previous study [56], we found an average of 10% DE transcripts were related mostly to fatty acid metabolism, structural components, and neuromuscular junction assembly.

### Lipid metabolism

FA lipids are used as an alternative energy source during fasting, starvation, and endurance exercise in oxidative muscles. Both So and Ta muscles contain oxidative myofibers, and both should express genes associated with FA metabolism (Fig 2). However, DE genes explicitly associated with FA transport and catabolism were increased in expression in So, which contains a higher percentage of oxidative fibers in its overall myofiber composition.

In contrast, the 269-fold increased expression of *Cyp4f39* in Ta suggests that lipid catabolism to generate ceramides is more active in Ta. Ceramide accumulation has been linked to insulin resistance in type II diabetes [57]. The increased *Cyp4f39* expression may allow Ta myofibers to utilize ceramides to shift from being insulin-sensitive facilitating efficient glucose uptake [26] to becoming insulin-resistant. Alternatively, increased expression of *Cyp4f39* and presumed increased ceramide levels in Ta might be associated with differential apoptosis, cell cycling, or autophagy.

### Glycogen and glycosaminoglycan metabolism

Glycogen acts as an energy reserve for myofibers by storing excess glucose molecules that are then utilized when blood glucose levels are low. Glycolytic or oxidative-glycolytic myofibers retain glycogen because they rely heavily on glucose as energy, and Ta has a high percentage of those myofibers; therefore, the increased expressions of the glycogen metabolic enzyme genes *Gys2, Pygm, Phka1, Phkg1*, and *Phkb* in Ta support the known metabolic composition of the muscle relative to So (Fig 3a; Table 2).

Hydrolysis of muscle glycogen to glucose occurs in lysosomes that engulf glycogen granules. The lysosome-associated DE genes, *Slc35d3, Atp6v0d2*, and *Galns*, exhibited increased expression in So, suggesting that lysosomal-related organelles in So and Ta myofibers perform slightly different functions (S1 Table). *Slc35d3* is associated with the biosynthesis of platelet-dense granules [58]. Platelet-dense granules contain high concentrations of calcium, adenine nucleotides, pyrophosphate, and polyphosphate molecules that enhance autophagy in lysosome-related organelles. *Atp6v0d2* is a part of vacuolar ATPases involved in proton translocation into vacuoles, lysosomes, or the Golgi apparatus to lower the pH [59]. The presence of *Atp6v0d2* suggested that lysosome-related organelles in So myofibers need to reduce their pH levels routinely. If lysosome-related organelles in So myofibers have enhanced autophagy, the pH levels within those organelles would fluctuate as the cell engulfs and metabolizes more molecules from the extracellular space and may require more proton pumps. *Galns* is involved in glycosaminoglycan biosynthesis. Glycosaminoglycans are cell surface proteins with branches of sugars projecting into the extracellular matrix to support cell identity, adhesion, and growth. *Galns* encodes for a protein that breaks keratan sulfate off of glycosaminoglycans [60] in lysosomes.

*Stab2, B3gat1*, and *Cspg5* are also involved in glycosaminoglycan synthesis, indicating that So and Ta synthesize distinct glycosaminoglycans associated with cell adhesion and proliferation (S1 Table, S2 Table). *Stab2* is involved in the endocytosis of metabolic waste products, including circulating hyaluronic acid (HA) that promotes cell proliferation once the plasma concentration decreases. *B3gat1* is involved in generating HNK-1 carbohydrate (CD57) cell surface epitope associated with cell adhesion [61]. *Cspg5* is involved in chondroitin sulfate synthesis, another molecule that can be added to proteoglycans and connected to cell adhesion, growth, migration, and receptor binding in the central nervous system (S1 Table). The high *Cspg5* expression in So suggests that neurons associated with the So muscle are marked differently.

### Glucose metabolism

The DE genes associated with glucose metabolism were increased in Ta more so than So. This imbalance seemed odd because both muscles use glycolysis. Yet *Eno2* and *Pck1* were increased in expression in So compared to the 11 isoenzymes in Ta, maybe due to the higher percentage of type II(B) glycolytic myofibers in Ta (Fig 4, Table 2).

Several of those isoenzymes are regulatory enzymes of glycolysis and gluconeogenesis. Glycolysis-related enzymes encoded by *Pfkm, Pkm Pfkfb3, Pfkfb1*, and *Pfkfb4* were increased in Ta (Fig 4b, Table 2). In contrast, two gluconeogenesis regulatory enzymes that control pyruvate’s reversion to glucose in skeletal muscle encoded by *Fbp2* and *Pck1* were expressed higher in Ta and So, respectively.

Gluconeogenesis in Ta might not be as tightly controlled across several steps as glycolysis seems to be on the 3-to-1 regulatory enzyme ratio. Gluconeogenesis may be controlled by the PFK2/FBPase2 complex instead of relying on the energy-costing enzyme PEP carboxylase as soleus seems to utilize. The gene *Pfkfb4* had a higher differential expression than *Pfkfb3. Pfkfb4* opposes *Pfkfb3* as they redirect glucose to the pentose phosphate pathway to promote detoxification of reactive oxygen species and lipid and nucleotide biosynthesis. *Fbp2*, which was also increased in expression in Ta, pulls F1,6-BP to convert back into fructose 6-phosphate. These findings combined may be indicating that Ta myofibers initiate gluconeogenesis at the PFK2/FBPase2 step of the glucose metabolism process to reduce energy loss.

### Contraction

Several components that contribute to the sarcomere formation were identified as DE between So and Ta, including MyHCs, myosin light chains, troponin subunits, and NMJ-associated proteins (Fig 5a-e).

The presence of type I (*Myh7*) and type II(B) (*Myh4*)-associated MyHC isoforms in the curated DE gene lists provided proof of concept for this analysis. The fold change in expression of *Myh7* and *Myh4*, 34.7- and 57.7-fold, respectively (Table 3), reflected the appropriate percentage of types I and II(B) myofibers in each muscle. So and Ta both have type II(A) and II(D) myofibers, so expressions of their specific MyHCs were predictably absent from our analysis. Our data also support a previous analysis correlating *Myh4* and *Myh7* with other genes associated with myofiber types I and II(B) [56], respectively (Table 3, S1 Table, S2 Table).

The myosin ELCs and RLCs correlated mostly to the appropriate muscle except for *Myl4* and *Myl6* (Fig 5c, Table 3). *Myl4* expression decreases in the skeletal muscle when mice reach embryonic day 15.5 during their development, and *Myl6b* is expressed in adult human slow-skeletal muscle, not in adult mice [62]. However, those genes were at a 3- and 54.1-fold increase in So over Ta, respectively (Fig 5c, Table 3).

The troponin complex subunits encoded by *Tnnt1, Tnni1*, and *Tnnc1* were unique and highly expressed in So (Fig 5b, Table 3), suggesting that troponin complex subunits in So might be modified relative to Ta to keep the actin filament in an open position longer, facilitating longer durations of contraction. The absence of unique subunits found in Ta could be interpreted as Ta possessing subunits that are also present in So. Our data support previously observed differences in troponin, tropomyosin, calsequestrin, myosin heavy chains, and myosin light chains between skeletal muscles [56].

Most of the NMJ-associated DE genes were identified among the genes with the largest fold-change and not through overrepresented pathway analysis, suggesting these genes may be underrepresented in curated pathway lists. Factors affecting NMJ specification and calcium handling have been theorized to be among the non-myofibrillar muscle-specific systems that allow for increased maximal isometric stress after P28 [5].

Synaptic modeling affects the motility or twitching of the skeletal muscle. Muscle identity is determined before innervation, yet innervation maintains muscle identity. For instance, during regeneration, only innervated So muscle can upregulate slow isoform mRNAs [63]. Yet, muscles do not completely change into another phenotype when neural cues are altered [64]. Intrinsic signaling from nerves and other sources, as well as environmental cues, dictate adult muscle phenotype [63].

Genes are associated with neurotransmission and synaptic vesicles [38] at the presynaptic side of NMJs in addition to clathrin-mediated endocytosis [34,37] and cell adhesion [36] of the postsynaptic side were highly expressed in So. The release and retrieval of the synaptic vesicles and their contents may play an important role in the slow-twitch mechanism. On the contrary, Ta had increased expression of genes associated with the presynaptic side of NMJs only, which modulate neuronal excitability [39], broaden the action potential [40], and slow down the degeneration of NMJs [41]. These genes may add to our understanding of how Ta and its associated neurons handle the stimulation necessary for rapid contraction. The DE genes related to the ion transport pathways may be associated with the postsynaptic side of NMJs in Ta myofibers, such as the voltage-dependent calcium channel subunits (S1 Table). However, all of this information and conjuncture is based on a small number of differentially expressed genes.

## Conclusion

So and Ta are characterized by unique metabolic and contractile properties. Approximately 10% of transcripts were differentially expressed between So and Ta. So and Ta were characterized with increased expression of genes involved in lipid metabolism and glucose and glycogen metabolism, respectively. So myofibers expressed unique isoforms of RLCs, ELCs, and troponin complex subunits, while genes encoding for other myofibrillar structures were expressed relatively similar between Ta and So myofibers. Gene expression of distinct components of non-myofibrillar structures such as NMJs suggests that electrochemical signaling could be different in So and Ta.

## Acknowledgments

The authors want to thank Arun Singh and Thai Tran for their assistance during this investigation’s data analysis stages.

## Supporting information

**S1 Fig. Evaluation of Reactome overrepresented pathway analysis**.

The statistically significant DE genes were filtered into curated molecular pathways using the *ReactomePA* R package. The Reactome overrepresented pathway analysis tool uses a provided list of genes and filters the genes into curated molecular pathways. Specific molecular pathways were identified as being over-represented based on the input genes exceeding the proportion of genes randomly expected for a particular molecular pathway. Genes with increased expression in So or Ta are labeled in red or blue, respectively. The heat map indicated the log2(FC) of 160 DE genes associated with the 16 overrepresented pathways identified by the Reactome database before filtering based on the absolute FC being greater than or equal to two. The sixteen pathways were combined into three major categories: *Contraction, Ion & Amino Acid Transport*, and *Glucose & Glycogen Metabolism* based on description similarities.

